# Noise-induced damage in the zebrafish inner ear endorgans: evidence for higher acoustic sensitivity of saccular and lagenar hair cells

**DOI:** 10.1101/2023.04.24.538073

**Authors:** Ieng Hou Lau, Raquel O. Vasconcelos

## Abstract

The three otolithic endorgans of the inner ear are known to be involved in sound detection in different teleost fishes, yet their relative roles for auditory-vestibular functions within the same species remain unclear. In zebrafish (*Danio rerio*), saccule and utricle are thought to play key functions in encoding auditory and vestibular information, respectively, but the biological function of lagena is not clear. We hypothesized that the saccule is the main auditory endorgan and lagena might serve an auditory function given its connectivity to the saccule and dominant vestibular function of the utricle in this species.

We investigated the acoustic sensitivity of the three otolithic endorgans in adult zebrafish by comparing the impact of acoustic trauma (continuous white noise at 168 dB for 24 h) on their sensory epithelia.

Noise treatment caused hair cell loss in both the saccule and lagena, but not in the utricle. This effect was identified immediately after acoustic treatment and did not increase 24h post trauma. Furthermore, hair cell loss was accompanied by a reduction in presynaptic activity measured based on Ribeye b expression but mainly in the saccule, supporting its main contribution for noise-induced hearing loss.

Our findings support the hypothesis that the saccule plays a major role in hearing and that lagena is also acoustically affected but with less sensitivity most likely extending the species hearing dynamic range.

**Summary statement:** Noise-treated zebrafish showed higher hair cell loss and synaptopathy in the inner ear saccule and, to some extent, in the lagena, confirming their higher sensitivity and contribution for hearing loss compared to utricle.

## Introduction

The inner ear of teleost fishes consists of three orthogonal semi-circular canals and three otolith endorgans. The otolithic endorgans, namely utricle, saccule, and lagena, consist of a sensory epithelium populated with hair cells that are overlaid by a gelatinous membrane and are mechanically coupled to a dense calcareous otolith (Lu and Popper, 1998). Movement of the sensory hair cells causes the excitation of afferent fibers along the axis of stimulation. Due to the difference in inertia between the sensory macula and the associated otolith, these endorgans function as biological accelerometers thus encoding linear acceleration, which can include orientation with respect to gravity, but also particle motion within the auditory range (Popper and Fay, 1999; Popper and Hawkins, 2018, 2021).

The sensory role of the inner ear endorgans can be vestibular and/or auditory, depending on the response frequencies and how the brain uses the information encoded in the VIIIth nerve afferents that project into the first order octaval nuclei in the brainstem (McCormick, 1999). The saccule has been described as the main auditory endorgan in teleost fishes (Brown et al., 2019; Coffin et al., 2012; Ladich and Schulz-Mirbach, 2016; Lu and DeSmidt, 2013; Smith et al., 2011), although the auditory sensitivity of utricle (Denton and Gray, 1979; Lu et al., 2004; Rogers and Sisneros, 2020) and lagena (Lu et al., 2003; Meyer et al., 2012; Sand, 1974; Vetter et al., 2019) has also been reported in a few species.

In earlier diverging vertebrates, such as birds and mammals, the inner ear endorgans typically show a well-defined segregation of their biological function (see Manley, 2000 for review). In fish, the extent to which the different endorgans serve an auditory versus vestibular role remains to be clarified. According to the ‘mixed function’ hypothesis, each otolithic endorgan serves both senses to varying extents (Platt and Popper, 1981; Popper and Fay, 1993; reviewed in Schulz-Mirbach et al., 2019, but this requires further evidence and might be species-specific. To date, the auditory contribution of single otolith endorgans has been tested only in a few fish species and the studies available mostly support the ‘mixed-function’ hypothesis (eg. goby *Dormitator latifrons* (Eleotridae) (Lu et al., 2010); midshipman/toadfish (Batrachoididae) (Fay and Edds-Walton, 1997; Sisneros, 2007; Vasconcelos et al., 2015); goldfish *Carassius auratus* (Cyprinidae) (Fay, 1984); and catfish *Ictalurus punctatus* (Ictaluridae) (Moeng and Popper, 1984). For instance, the midshipman fish *Porichthys notatus* has been well described regarding the auditory sensitivity of the inner ear endorgans and evidence shows that all three endorgans are sensitive to the frequency range of conspecific calls but with distinct auditory thresholds, thus probably complementing their function for acoustic communication and directional hearing (Sisneros, 2007, 2009). Given the vast morphological diversity in fish auditory systems, it is important that the hearing sensitivities are understood at the species level, and this requires that the contribution of each otolithic endorgan to hearing be evaluated.

The zebrafish *Danio rerio* is an otophysian teleost with Weberian ossicles connecting the inner ear to the swim bladder expanding the auditory sensitivity range. This species has become a well-established model organism in hearing research to investigate inner ear development, hair cell regeneration, and to test ototoxic agents and drugs treatments for auditory impairments (Bever and Fekete, 2002; Chiu et al., 2008; Ou et al., 2010; Varshney et al., 2016; Wang et al., 2015; Whitfield et al., 2002). The saccule and utricle of zebrafish are thought to play key functions in encoding auditory and vestibular information, respectively (Schulz-Mirbach et al., 2019; Whitfield et al., 2002). Removal of the utricle leads to impaired postural equilibrium, which does not occur when saccule is absent (Bianco et al., 2012; Moorman and Riley, 2000). Besides, larval zebrafish without utricle showed only slightly reduced auditory sensitivity at low frequencies <100 Hz, while manipulation of the saccule caused >30 dB hearing loss within 100–400 Hz during the first week post fertilization (Yao et al., 2016). In zebrafish, the lagena is the last to develop among the three endorgans, i.e. around 15 days post fertilization, and it is attached to the saccule on the posterior part at the adult stage (Bever and Fekete, 2002). In otophysian fishes, it is common that lagena is similar or even larger in size compared to the saccule, but its biological function remains unclear (Popper et al., 2003; Schulz-Mirbach and Ladich, 2016).

Based on the aforementioned, we hypothesized that the saccule is the main auditory endorgan and that lagena might serve an auditory function given its connectivity to the saccule and dominant vestibular function of the utricle in zebrafish. The major goal of this study was to investigate the impact of noise exposure on the hair cells and presynaptic function of the three otolithic endorgans in the adult zebrafish. The structural damage of noise trauma was compared between endorgans in order to evaluate their relative sensitivity to acoustic stimuli similarly to prior studies (Smith et al., 2011).

## Materials and Methods

### Animals

Wild type zebrafish (AB line) were originally obtained from China Zebrafish Resource Center (CZRC, China) and raised at the zebrafish facility of the University of Saint Joseph, Macao SAR. Fish were initially maintained in 10 L tanks in a standalone housing system (model AAB-074-AA-A, Yakos 65, Taiwan) with filtered and aerated water (pH balanced 7–8; 400–550 μS conductivity) at 28 ± 1 °C and under a 12:12 light: dark cycle. Specimens were fed twice a day with both dry food powder (Zeigler, PA, USA) and life artemia. A total of 44 adult zebrafish with 7-8 months old, 2.7-3.2 cm total length, and 0.180 – 0.240g body weight were used in this study.

All experimental procedures complied with the ethical guidelines regarding animal research and welfare enforced at the Institute of Science and Environment, University of Saint Joseph, and approved by the Division of Animal Control and Inspection of the Civic and Municipal Affairs Bureau of Macao (IACM), license AL017/DICV/SIS/2016.

### Acoustic treatments

Prior to noise exposure, all fish were transferred to 4L tanks for 7 days in a walk-in soundproof chamber (120a-3, IAC Acoustics, North Aurora, IL, USA) for isolation to reduce potential effects from captive noise (Lara and Vasconcelos, 2019). In these tanks, Sound Pressure Level (SPL) was around 103–108 dB re 1 μPa (LZS, RMS sound level measured with slow time and linear frequency weighting between 6.3–20 kHz) and conditions (light, temperature water quality, and feeding schedule) were maintained similar to stock.

After the isolation period, specimens were allocated randomly into three groups defined as “Control”, “Noise”, or “Noise+24h”, and transferred to the acoustic treatment glass tanks (dimensions: 50 cm length × 30 cm width × 25 cm height; 38 L), which were placed on top of a sponge base and two Styrofoam layers. The sound treatment was generated via an underwater speaker (UW30, Electro-Voice, MN, USA) positioned in the centre of the tank bottom on top of a sponge base that prevented direct contact with the glass. The speaker was connected to an amplifier (ST-50, Ai Shang Ke, China), which was connected to a laptop running Adobe Audition 3.0 for windows (Adobe Systems Inc., USA). In each trial, two specimens were treated at the same time by placing them inside a net box (dimensions: 15×15×15 cm, mesh size 1 mm) that was suspended underwater at 1 cm distance above the speaker (Fig. 1).

**Figure 1.**
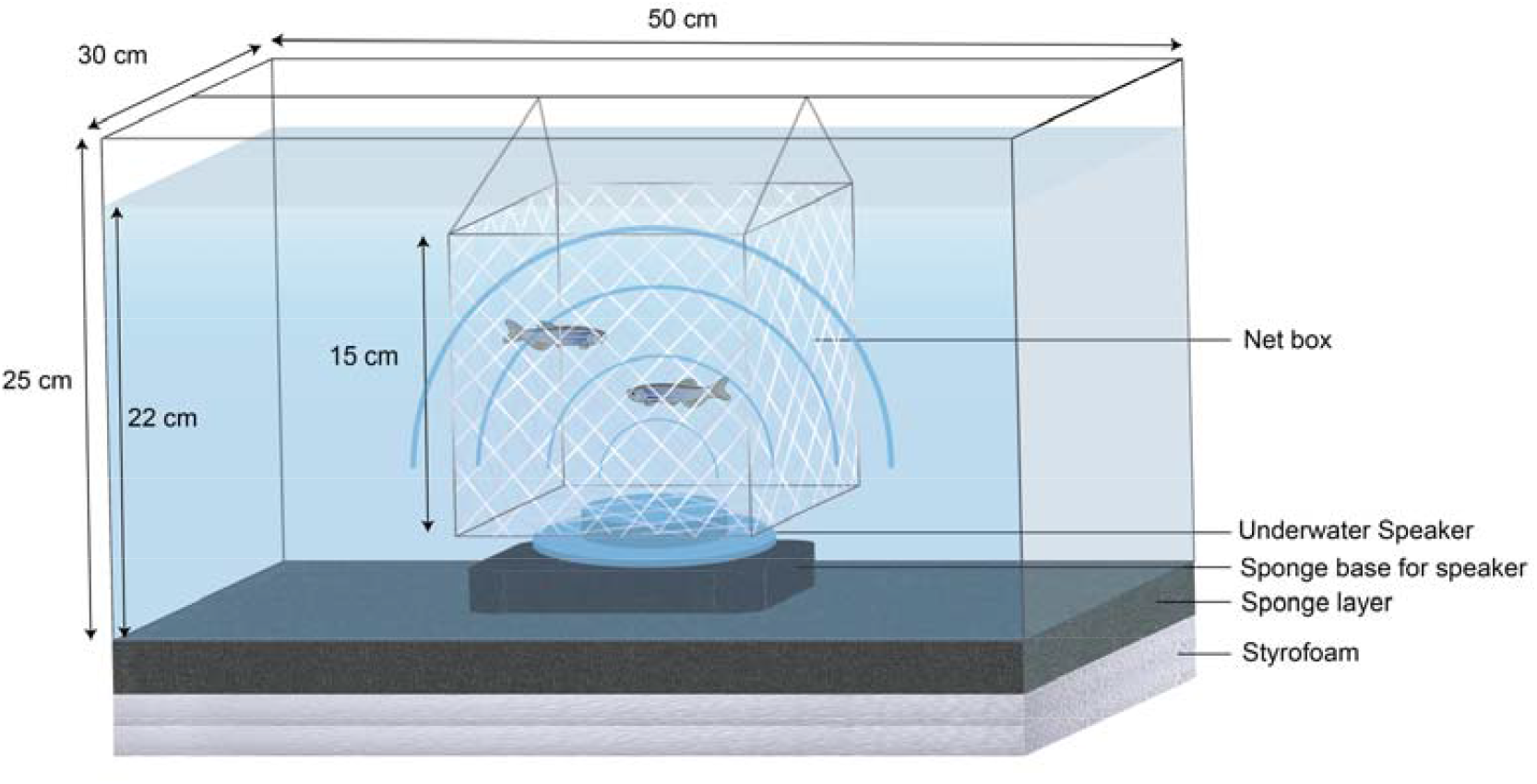
Acoustic treatment tank used to expose zebrafish inside a net box to sound playback (white noise at 168 dB re 1μPa) for 24 h.

The noise treatment consisted of white noise at 168 dB re 1 μPa (bandwidth: 100-1500 Hz) adjusted to the tank acoustic properties using Adobe Audition software tools to deliver a relative flat spectrum (Breitzler et al., 2020). Noise level was calibrated before each treatment so that the intended sound level (LZS) was reached at the center of the net box containing the fish. A SPL variation of ± 8 dB was registered inside the net box. In the “Control” treatment, the amplifier connected to the speaker was switched on but without playback and specimens were left in the tank for 24 h. In the “Noise” group, specimens were exposed to noise playback for 24 h. In “Noise+24h”, fish were firstly exposed to noise playback and then remained in the tank for additional 24 h under lab silent conditions. Sound recordings and SPL measurements were made with a hydrophone (Bruel & Kjær 8104, Naerum, Denmark; frequency range: 0.1 Hz to 120 kHz, sensitivity of – 205 dB re 1 V/μPa) connected to a hand-held sound level meter (Bruel & Kjær 2270).

The acoustic treatment was further calibrated with a tri-axial accelerometer (M20-040, frequency range 1**–**3 kHz, GeoSpectrum Technologies, NS, Canada) with the acoustic centre placed in the middle of the tank. The sound playback generated was about 140 dB re 1 m s^−2^, and most energy was in the vertical axis perpendicular to the speaker. Calculations were based on previously described methods using a MATLAB script paPAM (Nedelec et al., 2015).

At the end of the treatments, specimens were immediately euthanized with a MS-222 overdose (300 mg/L) buffered with sodium bicarbonate. Fish heads were subsequently removed and fixed overnight in 10% neutral buffered formalin solution (Sigma, USA), thoroughly rinsed with PBS during 5 min for at least 3 times, and finally stored in PBS at 4**°** C.

### Inner ear morphological analysis

We followed previously described procedures to extract zebrafish inner ear sensory epithelia from the three endorgans (Liang and Burgess, 2009). Sensory epithelia were dissected from both ears in each specimen under a stereomicroscope (Stemi 2000CS, Zeiss) and then immediately stained to quantify hair cell (HC) bundles and Ribeye b punctua based on previously described protocols (Wang et al., 2015). This protein is present in presynaptic ribbons that enhance neurotransmitter release and it is a measure of synaptic function (Nicolson, 2015).

Epithelia were firstly permeabilized with 1% Triton X-100 RT for 2 h, followed by blocking with 10% goat serum in PBS and incubated with primary antibody against ribeye b protein (mouse anti-zebra sh Ribeye b monoclonal antibody provided by T. Nicolson, Stanford University, CA, USA; 1:5000) at 4°C overnight. In the following day, samples were rinsed with 1% PBST for 2 hours, incubated with Alexa Fluor 647 IgG2a secondary antibody (Invitrogen, USA; 1:500) and Alexa Fluor 488 phalloidin (Invitrogen, USA; 1:500) for 2 h, rinsed for 30 min in PBS and subsequently stained for 1 min in DAPI solution. Finally, the samples were whole mounted with Fluoromount-G (Southern Biotech, USA) on glass slide, imaged with a confocal system (STELLARIS 5 LIA, Leica Microsystems CMS GmbH, German), and analysed with Leica Application Suite software X (LAS X, Leica Microsystems Leider Lane) and image J (Version 1.53e, National Institutes of Health, USA).

Hair cell bundles and nuclei were quantified in different squared regions of 900 mm^2^ across the sensory epithelia of the three endorgans (Fig. 2). Ribeye b puncta were also quantified in the same regions with 1–2 additional regions on the edge of the saccule and lagena, where higher amount of puncta were typically found. The epithelial regions were selected based on Wang et al. (2015) and Coffin et al. (2012). For saccule, they were located at 5%, 25%, 37.5%, 50%, and 75% across the length of the rostral-caudal axis. In the lagena, the regions were roughly at 25%, 50%, 75% and 95%, across the rostral-caudal axis. For the utricle, four regions were selected based on a rectangular projection (see Fig. 2). All HC bundles, nuclei and ribeye b puncta within or overlapping the square outlines were included in the counts. The two inner ears of each fish were analysed and the data of each endorgan was averaged, therefore only one value per specimen was included in the analysis.

**Figure 2.**
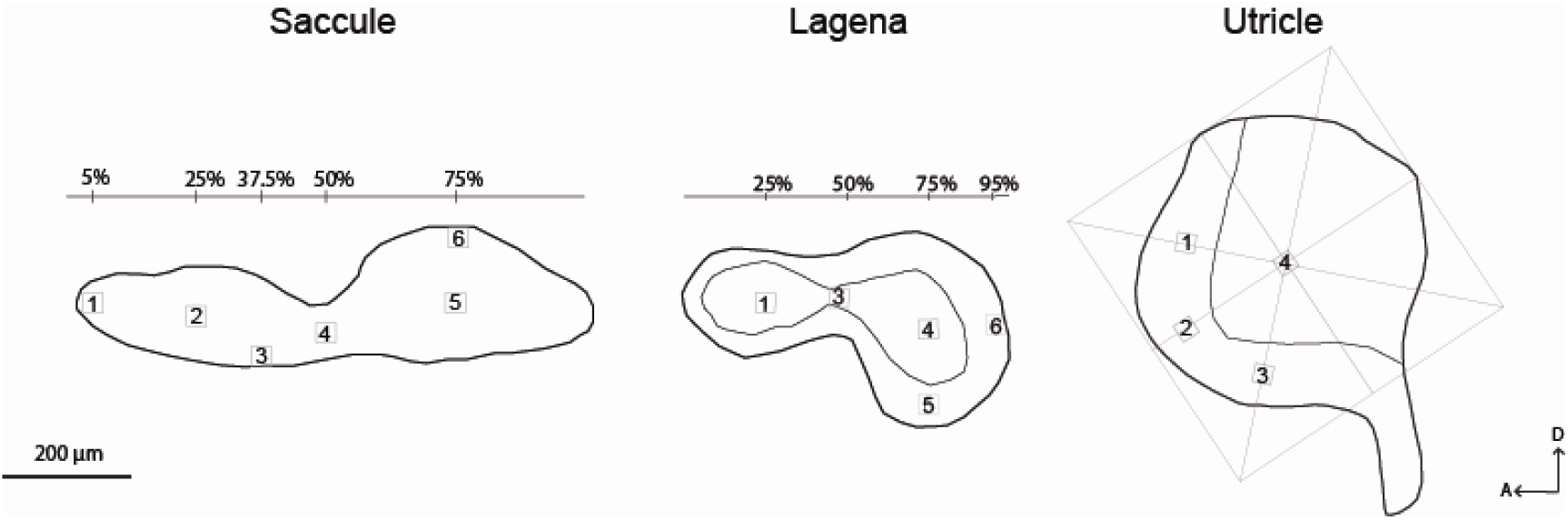
Diagram of representative inner ear endorgans from a zebrafish adult showing the squared regions (900 μm^2^) defined for hair cell bundle and Ribeye B puncta quantification. D, dorsal; R, rostral.

### Statistical Analysis

The differences in HC bundles and Ribeye b puncta between experimental groups were tested with two-way ANOVA with acoustic treatment and epithelial regions as independent factors, and then followed by Fisher’s LSD post hoc multiple comparisons to detect specific differences between treatments within each region. The relationship between HC bundles and the nuclei was determined with Pearson correlation and fitted with a regression line.

All assumptions for parametric analyses were confirmed through the inspection of normal probability plots and by performing the Levene’s test for homogeneity of variances. All statistical tests and graphics were performed using Matlab (MathWorks, Inc., USA) and Prism 9 (GraphPad, USA).

## Results

Noise exposure for 24 h caused significant HC loss in the saccule (F_2, 162_ = 48.09, P<0.001, Fig. 3A, C) for all epithelial regions analysed except the S4 region, with a 36-59% reduction in HC bundles (S1: P< 0.001; S2: P< 0.001; S3: P< 0.001; S5: p < 0.001, Fig. 3, 4). This HC bundle loss did not vary with the epithelial location (F_8, 162_ = 1.206, P>0.05), meaning that the acoustic trauma affected similarly the whole saccule.

**Figure 3.**
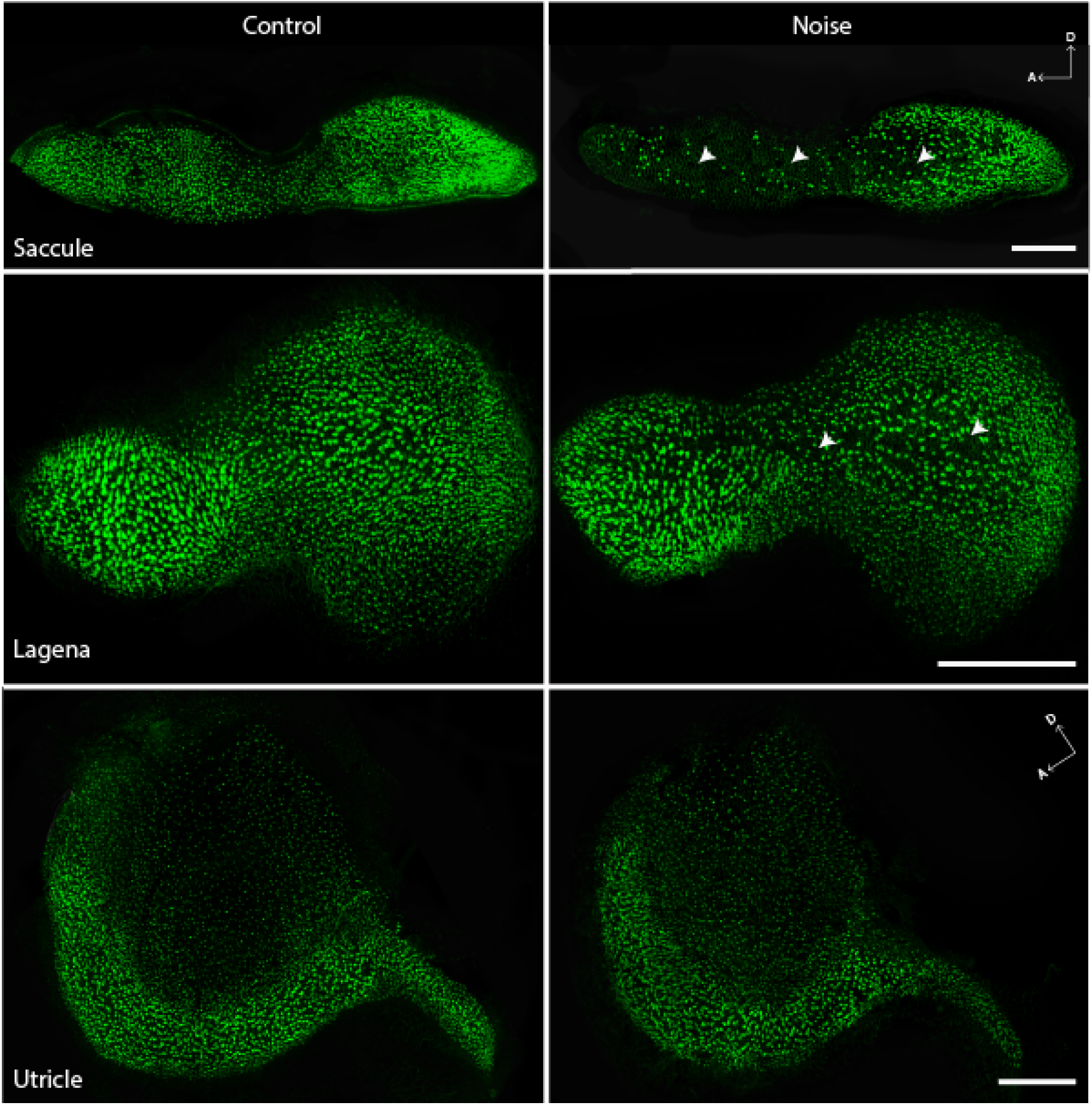
Representative images of the entire sensory epithelia of the three inner ear endorgans (saccule, lagena and utricle) of zebrafish adults reconstructed with the 3D function of LAS X software (Leica Microsystems) showing Phalloidin-stained hair cells from the control and noise-treated group. Noise-induced hair cell bundle loss is clearly shown for the saccule and to some extent in lagena. Utricular epithelia did not reveal noise-induced morphological changes.

**Figure 4.**
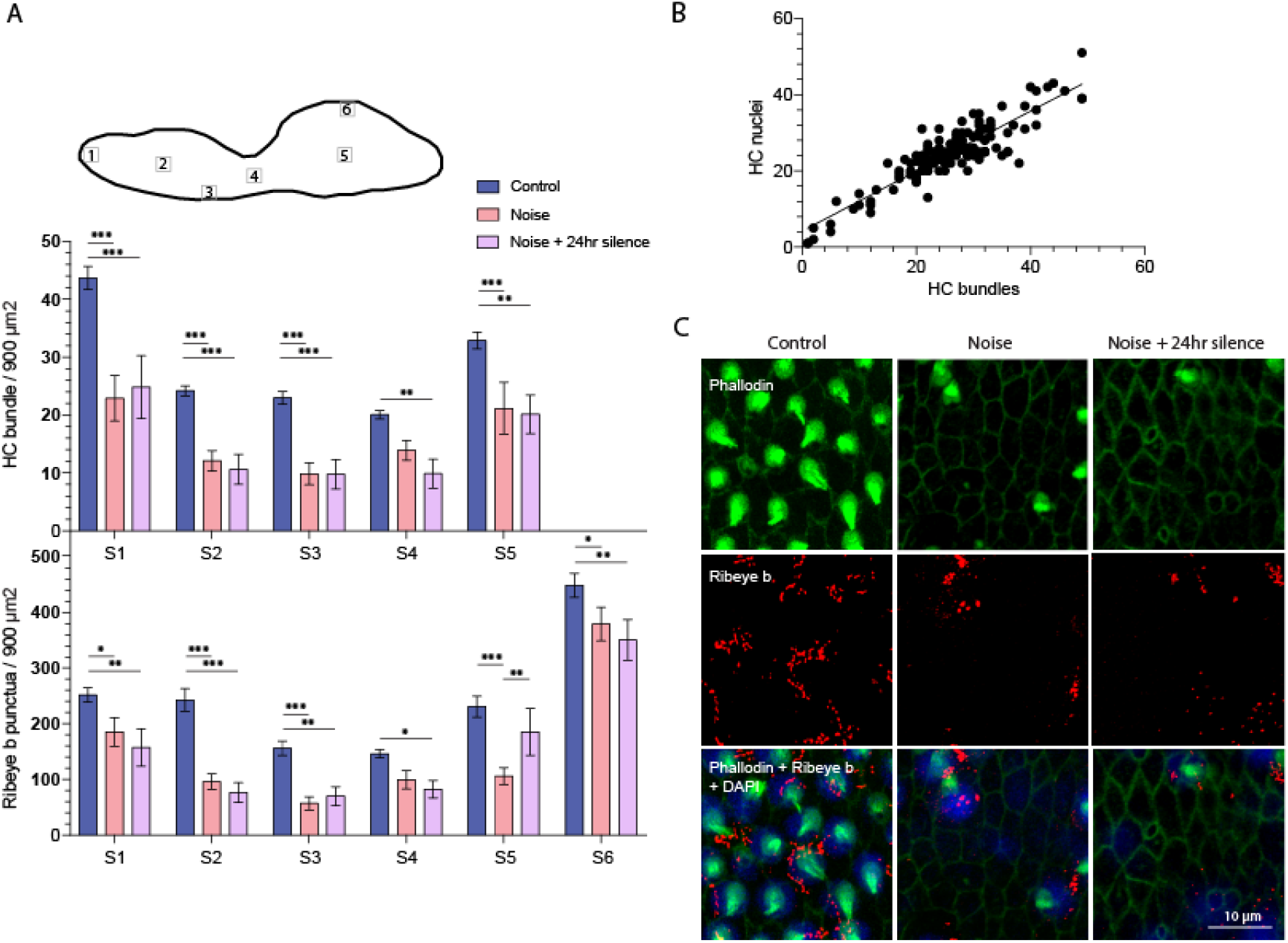
A) Variation in hair cell (HC) bundles and Ribeye b punctua in zebrafish saccular epithelia under different experimental conditions -control, noise treatment (24h), and noise treatment followed by 24h silent period. A diagram of the saccule is shown to indicate the epithelial regions analyzed. Values are mean ± SE. HC: F_2, 195_ = 43.87, P< 0.001; Ribeye b: F_2, 162_= 48.09, P< 0.001. Post hoc tests * P< 0.05, ** P< 0.01, *** P< 0.001. B) Correlation between HC nuclei and HC bundles (r (126) = 0.8971, P< 0.0001) fitted with a regression line (Y= 0.7876X + 4.179, R^2^ = 0.8048). C) Representative images of saccular epithelia (from S2 region) showing Phalloidin-stained HCs (green), presynaptic Ribeye b puncta (red), and DAPI-stained HC nuclei (blue) for the three experimental groups.

Noise-induced morphological changes were accompanied by reduced pre-synaptic activity quantified based on Ribeye B expression (F_2,195_ = 43.87, P<0.001) for most epithelial regions analysed (S1: P= 0.019; S2: P< 0.001; S3: P< 0.001; S5: P< 0.001; S6: P= 0.010, Fig. 4) with an overall 22-69 % decrease in protein expression. Such change varied with the saccular epithelial region (F_10,195_= 1.927, p= 0.044). In this case, a more significant decrease (55-69%) was registered in the anterior saccular epithelium (S2 and S3), and no significant difference was found in the S4 region.

In general, additional 24 h period post -trauma neither caused changes in saccular HC (F_1,99_= 0.2056, P> 0.05) nor in Ribeye B expression (F_1,120_= 0.0001, P> 0.05). Yet, the region S5 located in the middle of the posterior saccular epithelia revealed slight increase in Ribeye b (P= 0.010).

Moreover, the amount of HC bundles was significantly correlated with HC nuclei in the saccule (r_126_= 0.8971, p< 0.0001), Fig. 4B, C). Noise-induced HC bundle loss was associated with lower amount of HC nuclei, suggesting that acoustic trauma caused cell death rather than only stereocilia damage and/or bundle loss (r (126) = 0.8971, p<0.0001).

The acoustic treatment also induced significant HC loss in the lagena (F_2,,106_= 10.44, P< 0.001, Fig. 3, 5), specifically in the regions L1, L2 and L3 that were located in the anterior, middle and posterior epithelia, respectively (L1: P= 0.020; L2: P= 0.003; L3: P= 0.016, Fig. 5). Noise exposure caused about 21-23 % of HC bundle loss in these regions. Such cellular damage was accompanied by significant changes in Ribeye b expression (F_2,169_= 3.235, P= 0.042), but significant differences were only found in L3 (p= 0.010), as shown in Fig. 5A, B.

**Figure 5.**
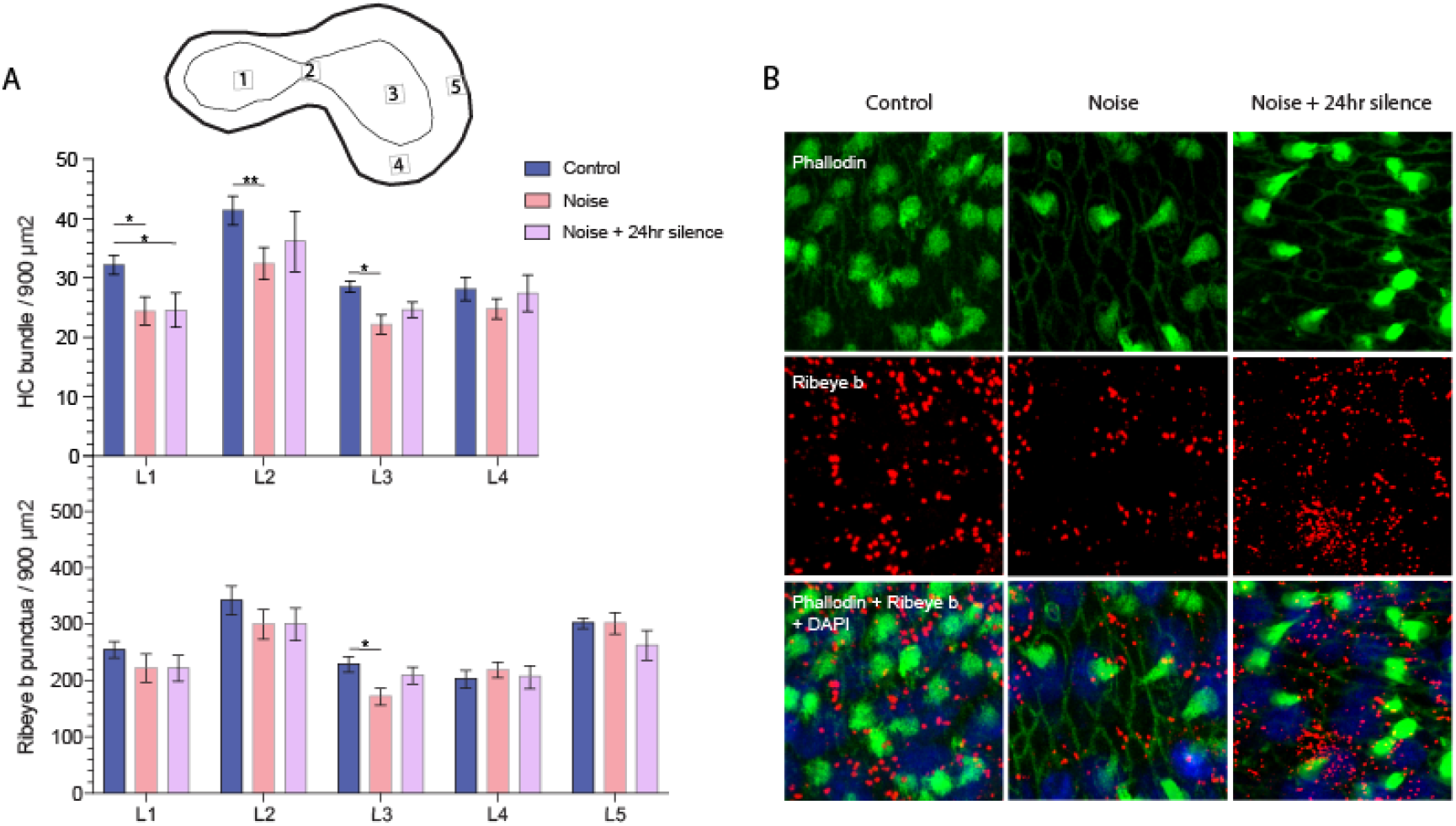
A) Variation in hair cell (HC) bundle and Ribeye b punctua in zebrafish lagena under different experimental conditions - control, noise treatment (24h), and noise treatment (24) followed by additional 24h silent period. A diagram of lagena is shown with the epithelial regions analyzed. Values are mean ± SE. HC: F_2,,106_= 10.44, P< 0.001; Ribeye b: F_2,169_= 3.235, P= 0.042. Post hoc tests *P < 0.05, **P < 0.01, ***P<0.001. C) Representative images of lagena epithelia (from L3 region) showing Phalloidin-stained HCs (green), presynaptic Ribeye b puncta (red), and DAPI-stained HC nuclei (blue) for the three experimental groups.

Finally, the utricle was not significantly affected by the acoustic treatment, neither in terms of HC loss (F_2,79_= 2.270, P> 0.05) nor in presynaptic activity (F_2,80_= 3.069, P > 0.05) (Fig. 3, 6).

**Figure 6.**
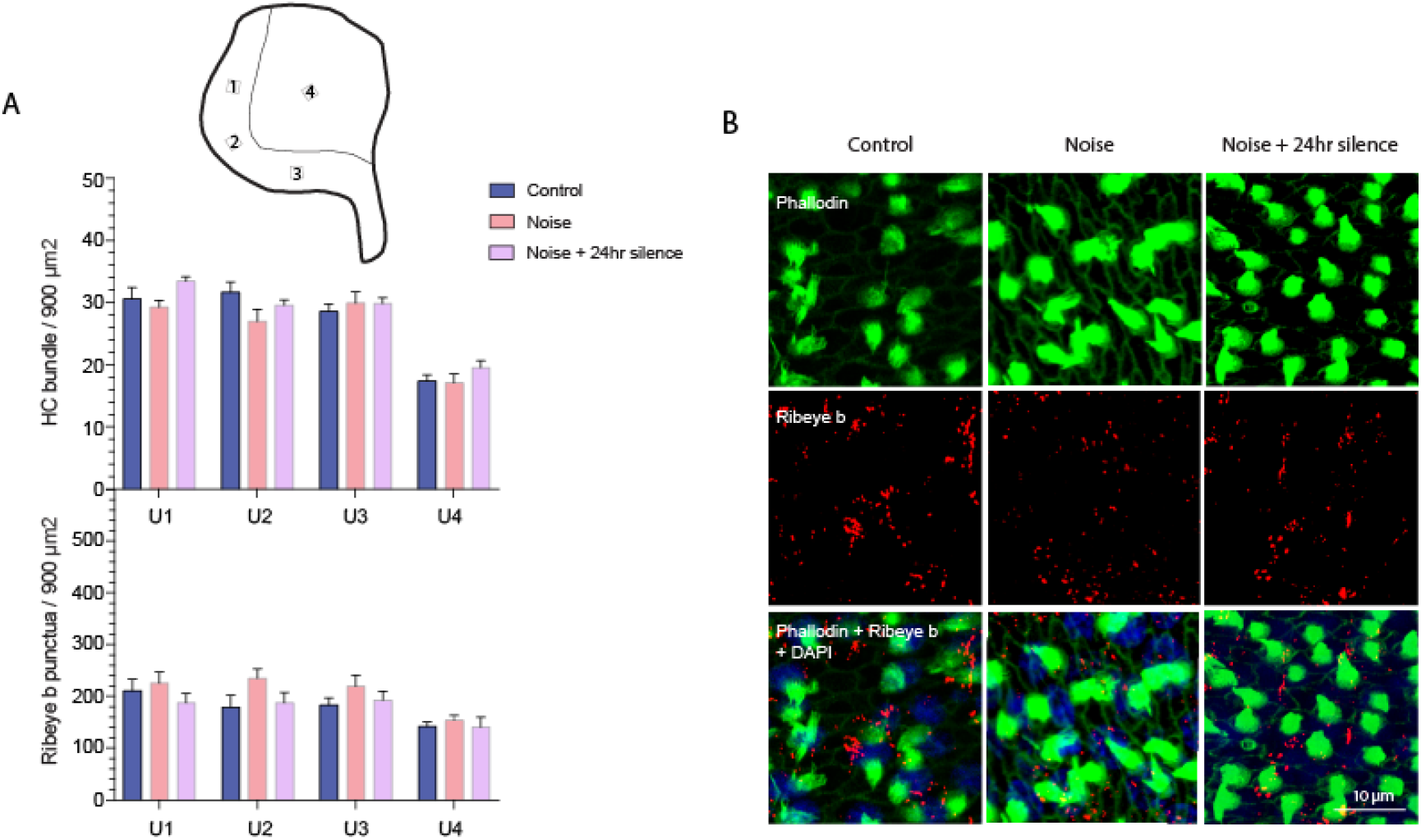
A) Variation in utricular hair cell (HC) bundles and Ribeye b punctua in experimental groups - control, noise treatment (24h), and noise treatment (24) followed by additional 24h silent period. A diagram of utricle and specific epithelial regions analyzed are indicated. Values are mean ± SE. HC: F_2, 79_= 2.27, P > 0.05; Ribeye b: F_2, 79_= 2.97, P > 0.05. C) Representative images from utricular epithelia (U2 region) showing Phalloidin-stained HCs (green), presynaptic Ribeye b puncta (red), and DAPI-stained HC nuclei (blue) for the three experimental groups.

Overall comparison of noise-induced HL loss and Ribeye b confirmed the differences between the endorgans (HC: _F1,46_ = 28.10, P<0.0001; Ribeye B: F_1,39_ = 5.409, P = 0.025) – Fig 7.

**Figure 7.**
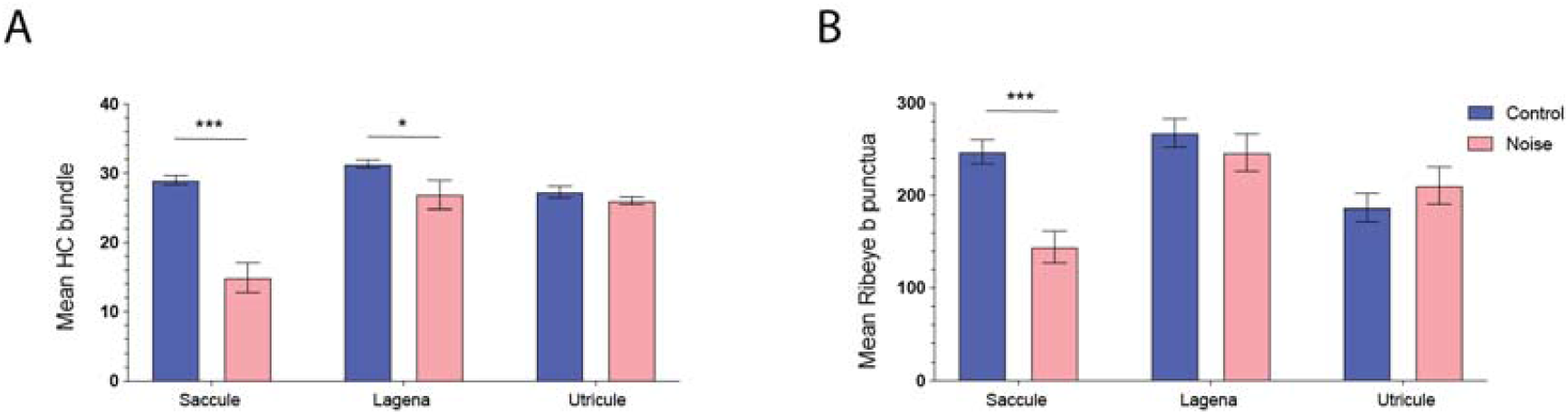
Comparison of mean hair cell (HC) bundles (A) and Ribeye b punctua (B) between the inner ear endorgans of zebrafish after noise treatment and silent control conditions. Values are mean ± SE. HC: _F1,46_ = 28.10, P<0.0001; Ribeye B: F_1,39_ = 5.409, P = 0.025. Post hoc tests *P < 0.05, **P < 0.01, ***P<0.001.

## Discussion

The present work compared the acoustic sensitivity between the three otolithic endorgans of the zebrafish inner ear based on their structural-functional changes after acoustic trauma. Our results showed the highest effects at the morphological and synaptic level in the saccule. Lagena was also acoustically affected but with less damage, contrasting with the utricle that did not reveal noise-induced epithelial changes at the noise level investigated (168 dB re 1μPa for 24 h). We provide evidence that the zebrafish inner ear endorgans may have different auditory sensitivities most likely to extend the species hearing dynamic range and to complement their differential roles on auditory-vestibular function.

### Noise-induced hair cell loss

The impact of noise exposure on the fish inner ear has been addressed in a few studies. Most researchers focused on the saccule, which is considered to play a major role in hearing in this taxon (see Ladich and Schulz-Mirbach, 2016; Schulz-Mirbach et al., 2019 for review), and reported evidence of a tonotopic organization of the saccular epithelia with the rostral region more sensitive to high-frequency noise and the caudal region affected by low frequencies (Breitzler et al., 2020; Enger, 1981; Han et al., 2022; Schuck and Smith, 2009; Smith et al., 2006; Smith et al., 2011).

In the current work, continuous exposure to white noise at 168 dB for 24 h caused a significant reduction in hair cell bundles throughout the whole saccule. Our results did not reveal a significant variation in hair cell loss with the epithelia region, but we found a higher impact in the rostral region of the saccule (48-57%) compared to the central and caudal regions (< 36%). This pattern is similar to prior studies that investigated the same species after white noise exposure. In our previous study, Breitzler et al. (2020) exposed zebrafish to white noise at 150 dB re 1 μPa for 24 h and identified significant hair cell loss (up to 27%) mostly in the rostral region of the saccule. More recently, (Han et al., 2022) treated zebrafish with white noise generated by a woofer air speaker at 140 dB re 1 μPa for 6 h and reported up to 22% decrease in saccular hair cells also in the rostral region. Both studies found the highest auditory thresholds shifts at 1000 Hz (Han et al., 2022) and 1000-2000 Hz (Breitzler et al., 2020), and such high frequencies are typically detected in the rostral saccular region. These frequencies fall within the best hearing range of the species (Breitzler et al., 2020; Han et al., 2022; Monroe et al., 2016), and it is likely that a white noise exposure may induce higher damage in the epithelial regions that detect the most sensitive frequencies. The higher amplitude used in our study (168 dB re 1μPa) is probably the cause for the higher and extended hair cell loss detected compared to Han et al. (2022) and Breitzler et al. (2020).

In the present study, we also evaluated the impact of noise on the lagena and utricle. We found about 21–23 % hair cell loss in the lagena in three different epithelial regions across the anterior-posterior axis, contrary to the utricle that did not reveal any cellular damage. This is one of the first reports comparing the impact of noise among the three inner ear endorgans in a fish species. Han et al. (2022) investigated the effect of 140 dB white noise for 6 h on zebrafish and identified over 40% hair cell loss in the caudal epithelia of the lagena and up to 25% hair cell loss in both saccule and utricle. The results obtained by these authors are difficult to compare with the present work since they used a very different treatment setup with a woofer air speaker and no information is provided in particle motion. Besides, the authors identified higher noise-induced damage in the lagena compared to the saccule, which is difficult to interpret. Our findings agree with the work by Smith et al. (2006) that exposed goldfish to white noise (170 dB re 1 μPa RMS) for 48 h and identified highest apoptotic cells in the saccule immediately following noise exposure but also in the lagena 24h post treatment. According to this study, the utricle that did not reveal any noise-induced apoptotic cells. Furthermore, Wang et al. (2019) reported an increase in apoptosis and cell proliferation in the zebrafish inner ear after a blast wave exposure that caused visceral hemorrhage. The saccule and lagena were the most affected endorgans with both around 400 apoptotic labeled cells, contrasting with the utricle that only revealed about 60 dead cells.

We have analysed noise-induced changes immediately after acoustic treatment and 24 h post treatment. The results showed that such additional period neither caused changes in saccular hair cells nor in presynaptic activity. Apart from the primary auditory hair cell death that occurs during noise exposure, secondary cellular death can be induced post trauma due to reactive oxygen and nitrogen species (Yamashita et al., 2004). Nevertheless, in this study no further HC bundle loss was found on the saccule one day after noise exposure, similar to the result reported by Schuck and Smith (2009). This might be due to different time delays in ROS and RNS formation, or because the intense damage on the saccule masked the effect of such processes.

### Noise-induced synaptopathy

The present work also identified significant noise-induced reduction in presynaptic activity based on Ribeye b quantification mostly in the saccule. The lagena also showed a slight reduction but only in one epithelial region. Ribeye b is known to be present in presynaptic ribbons that promote fast signal transmission between the auditory receptors and postsynaptic neurons, thus being critical in sound encoding (Kindt and Sheets, 2018; Nicolson, 2015). A reduction in Ribeye b protein has been associated to synaptopathy in hair cells after acoustic trauma mainly in mammals (Fernandez et al., 2015; Liberman and Kujawa, 2017; Valero et al., 2017) but also in fish (Uribe et al., 2018; Wang et al., 2015; Wong et al., 2022), leading to auditory threshold shifts and Noise-Induced Hearing Loss (NIHL) (Kindt and Sheets, 2018; Wong et al., 2022).

In zebrafish, a previous study reported a decrease in Ribeye b along with hair cell loss in the saccule associated to increasing size and ageing (Wang et al., 2015). Other researchers have also identified a reduction in the amount of Ribeye b ribbons in the neuromasts of noise-treated larval zebrafish (Holmgren and Sheets, 2021; Uribe et al., 2018). Recently, our lab (Wong et al., 2022) exposed zebrafish adults to white noise (150 dB re 1 μPa) of varying temporal patterns during 24 h and found significant Ribeye b reduction in the saccule mostly with the continuous noise (up to 40–50%) compared to intermittent regimes. In the present study, we found up to 69% reduction in protein expression, probably due to the higher noise amplitude used.

Our findings support the hypothesis that the saccule plays a major contribution for noise-induced hearing loss and that the sensory function of lagena is also acoustically affected but to a less degree.

### Differential role of the inner ear otolithic endorgans

According to our results, the saccule was the most sensitive endorgan to noise exposure based on the higher saccular hair cell loss and synaptopathy observed. These findings agree with most literature on fish auditory systems that report saccule as the major auditory endorgan (Brown et al., 2019; Coffin et al., 2012; Ladich and Schulz-Mirbach, 2016), including in zebrafish (Brown et al., 2019; Coffin et al., 2012; Ladich and Schulz-Mirbach, 2016; Lu and DeSmidt, 2013; Smith et al., 2011). Fish species that have accessory hearing structures and improved auditory abilities, such as otophysans that include zebrafish, possess Weberian ossicles connecting the saccule to the swim bladder, which is typically correlated with higher auditory sensitivities and/or expanded hearing range (reviewed in Braun and Grande, 2008). Otolith-removal experiments (Lu and Xu, 2002) and single-unit or field potential recordings from saccular hair cells (Lu et al., 1998; Lu and Xu, 2002; Vasconcelos et al., 2015; Yao et al., 2016) also provided evidence that the saccule plays a major role in sound detection. Since field potential recordings from the zebrafish endorgans would be very difficult to perform due to the difficult access to the inner ear compared to other fish models (eg. midshipman fish, toadfish, and gobies (Brown et al., 2019; Lu et al., 2010; Vasconcelos et al., 2015), the present work provides an essential confirmation of the major sensitivity of the saccule to acoustic stimuli in the adult zebrafish.

We further demonstrated that lagena was also acoustically affected but comparatively less, in terms of hair cell loss and reduction in presynaptic activity. Lower noise-induced effects in the lagena suggest higher auditory thresholds, which implies that such endorgan probably aids fish in sound detection, especially at high sound levels close to the sound source when saccule becomes saturated. This extended hearing dynamic range would occur mostly in the vertical plane, since the hair cells of both lagena and saccule are typically oriented along this plane (Schulz-Mirbach et al., 2019). Up to date, the functional role of lagena is not clear and likely varies among different fish species due to the high diversity in morphology and connection/proximity to the swimbladder (Schulz-Mirbach and Ladich, 2016). Evidence of auditory function of lagena was reported in only a few studies. For example, Vetter et al. (2019) showed that lagena of the midshipman *P. notatus* was sensitive to low frequencies but with higher thresholds compared to the saccule. Other studies focusing on lagenar function (either microphonics or afferent recordings) also demonstrated its acoustic sensitivity and suggested an important role for directional hearing in the vertical plane (Fay, 1984; Lu et al., 2003; Sand, 1974).

Finally, the present work showed the absence of hair cell loss and synaptopathy in the utricle. This endorgan is known to be mainly involved in detecting gravitational forces for vestibular sensing (Riley and Moorman, 2000; Roberts et al., 2017). Nevertheless, a few studies support the idea that utricle may serve an auditory role to some extent (Lu et al., 2004; Rogers and Sisneros, 2020), including in larval zebrafish (Yao et al., 2016).

To date, the auditory contribution of single otolith endorgans has been tested only in a few fish species (Fay, 1984; Fay and Edds-Walton, 1997; Lu et al., 2010; Moeng and Popper, 1984; Sisneros, 2007; Vasconcelos et al., 2015) and such studies mostly support the ‘mixed-function’ hypothesis. Our results agree with this hypothesis, providing evidence that saccule is the major auditory endorgan and lagena is also acoustically sensitive. The utricle was not affected by the acoustic stimuli tested, most likely because of its dominant vestibular function (Schulz-Mirbach et al., 2019; Whitfield et al., 2002).

Given the vast diversity in structure and function of fish auditory systems, it is important that future research considers distinct taxonomic groups, as well as the contribution of each otolithic endorgan for hearing within each species and their potential differences in sensitivity to acoustic trauma.

## Acknowledgments

We thank Dr. Teresa Nicolson (Stanford University, CA, USA) and Dr. Jiping Wang (Affiliated Sixth People’s Hospital of Shanghai Jiao Tong University, Otolaryngology Institute of Shanghai Jiao Tong University, Shanghai, China) for providing the Ribeye b antibody.

This study was supported by the Science and Technology Development Fund (FDCT), Macao (ref. 046/2018/A2 and 0068/2020/A2).

## Authors’ contribution statement

R.O.V. conceived the idea, designed and supervised the research, and obtained funding. I.H.L. carried out the experiments, conducted data acquisition and analysis. Both authors contributed to critical interpretation of results and wrote the manuscript.

